# In situ community dynamics influences the temperature- and light- dependent succession of seasonal phytoplankton

**DOI:** 10.1101/2021.03.03.433693

**Authors:** Margot Tragin, Stefan Lambert, Jean-Claude Lozano, François-Yves Bouget

## Abstract

Temperature and light play a crucial role in regulating phytoplankton blooms in the Ocean. To assess the importance of these two parameters experimentally, microcosms were conducted on seven picoplankton communities (<3 μm) sampled in December, March, June and September 2015 and 2016 in the North Western Mediterranean Sea. Each community was exposed to 4 realistic seasonal conditions (December, March, June and September). Metabarcoding was used to investigate the eukaryotic diversity in the 56 microcosms experiments in parallel to high-frequency monitoring of environmental diversity in the sea. The three major lineages identified were the Stramenopiles, Alveolata and Archaeplastida. Overall, the five-day incubations were not sufficient to reshape the initial microbial communities completely. The microcosm outcome was strongly influenced by the dynamics of phytoplankton starting communities. In pre-bloom conditions, phytoplanktonic species were the most sensitive to temperature and light conditions. During a bloom, species belonging to diatoms or Chlorodendrophyceae usually did not respond to light and temperature in microcosms and continued to bloom independently of the applied seasonal condition. Together, these results suggest that light and temperature seasonal conditions play a crucial role in regulating phytoplankton dynamics in pre-bloom conditions and biotic interactions may be preponderant in bloom and post-bloom conditions.

## Introduction

As soon as marine planktonic life was discovered in the late XIX century, several studies reported phenology of plankton abundance [1]. The organisms size, whose seasonal pattern were described, evolved together with improvements of detection methods, from light microscopy to molecular techniques. The annual temporality of zooplankton such as copepods and large size phytoplankton such as diatoms and dinoflagellates has been studied for over a century in Northern Mediterranean Sea waters [1–5]. Later, small size picophytoplankton (cell size < 3 μm), such as the green picoalga *Micromonas* sp. were found to seasonally grow also in the Mediterranean sea [6, 7]. In the nineties, extensive microscopy-based taxonomic survey reported that photosynthetic dinoflagellates and diatoms had strong seasonal patterns in NW Mediterranean Sea coastal waters and described sequential phytoplankton spring blooms in the bay of Villefranche sur Mer (France): diatoms blooming in late spring and autumn while dinoflagellates being more abundant in summer [8]. More recently, the development of molecular analyses and high throughput sequencing enabled the tracking of all planktonic organisms’ temporal distribution, including the picoplankton [9]. In recent years, the development of molecular analyses and high throughput sequencing allowed the extensive monitoring at an unprecedented taxonomic level, the temporal succession of all planktonic organisms including the smallest protists (i.e., picoplankton cell size < 3 μm).

A time series metabarcoding approach conducted in the bay of Banyuls sur Mer (North Western Mediterranean Sea, France) over seven years revealed a strong temporal factor driving the biogeography of dinoflagellates and pico-sized green alga (Chlorophyta) belonging to the Mamiellophyceae class (*Bathycoccus* and *Micromonas*) [9, 10]. A quarter of photosynthetic reads were commonly assigned to Chlorophyta using 18S rRNA metabarcoding, describing pan-oceanic datasets [11]. The beginning of bloom events appears to be predictable by mathematical models that combine both nutrients, temperature and light parameters with hydrodynamics factors [12]. *In situ* high frequency sampling also unveiled the importance of daily variations in wind or rain on microbial community structures [13]. In addition to meteorological events, hydrographic modifications of environmental factors, which are thought to drive the seasonality of plankton, biotic interactions such as parasitism [14] were assumed to be influencing the end of blooming dynamics in marine waters [15].

Microcosm experiments have been used for decades to address a large panel of questions experimentally. Methodological approaches are not standardized regarding parameters such as temperature, nutrients or pH and targeted organisms (i.e., prokaryotes, pico- or nanoeukaryotes, phototrophs or heterotrophs). The duration of incubations commonly varied from several hours [16], to days [17], or even weeks [18, 19]. In the bay of Banyuls, two parameters, temperature and daylight, account for half of eukaryotic phytoplankton variability in one of the most extended metabarcoding time series (7 years) reported to date [9]. In this study, we combine microcosms and time series approaches in the bay of Banyuls to investigate the resilience of protists seasonality to light and temperature parameters, set up to mimic realistic seasonal conditions of daylight (photoperiod, intensity) and temperature. Over five days, incubations were performed that matched the gap between two successive samples in our high frequency (159 samples between 2015-2017) time series. These experiments aim at addressing the following questions: (1) How do abrupt changes in light and temperature alter picoeukaryotic communities? (2) How quickly can we change the protist communities from the initial season towards protist communities representative of the mimicked season? (3) How does the starting community dynamics influence microcosm outcome?

## Material and Methods

### Microcosms experiments

Five liters of seawater were sampled at SOLA buoy in the bay of Banyuls (42°31′N, 03°11′E, France). Three μm filtration was used to remove large grazers and nanoplanktonic predators such as ciliates and flagellates since they can induce strong bias in small volume microcosms. Four times 400 ml pre-filtered seawater were incubated in 500 ml aerated culture flasks (Sarstedt) in homemade incubators equipped with white wide spectrum LEDs mimicking realistic light conditions (photoperiod and light intensity, Fig. 1). For each date, microcosms were conducted in triplicates. In addition, the complete set of 84 incubations (7 dates x 4 conditions x triplicates) was done twice with and without nitrate and phosphate enrichments (NP). Five μM of NaNO_3_ (SIGMA Aldrich) and 0.2 μM of Na_2_HPO_4_ (SIGMA Aldrich) were added in NP enriched microcosms, which corresponded to maximal annual concentrations measured in the Banyuls Bay [9]. The planktonic abundance in the 56 microcosms (x3) was followed daily using an ACCURI C6 flow cytometer (BD Biosciences) as previously described [9]. After five days of incubation, triplicates were pooled, filtered on 0.22 μm Sterivex (Merck-Millipore) and stored at −80 °C prior to DNA extraction.

**Fig. 1:**
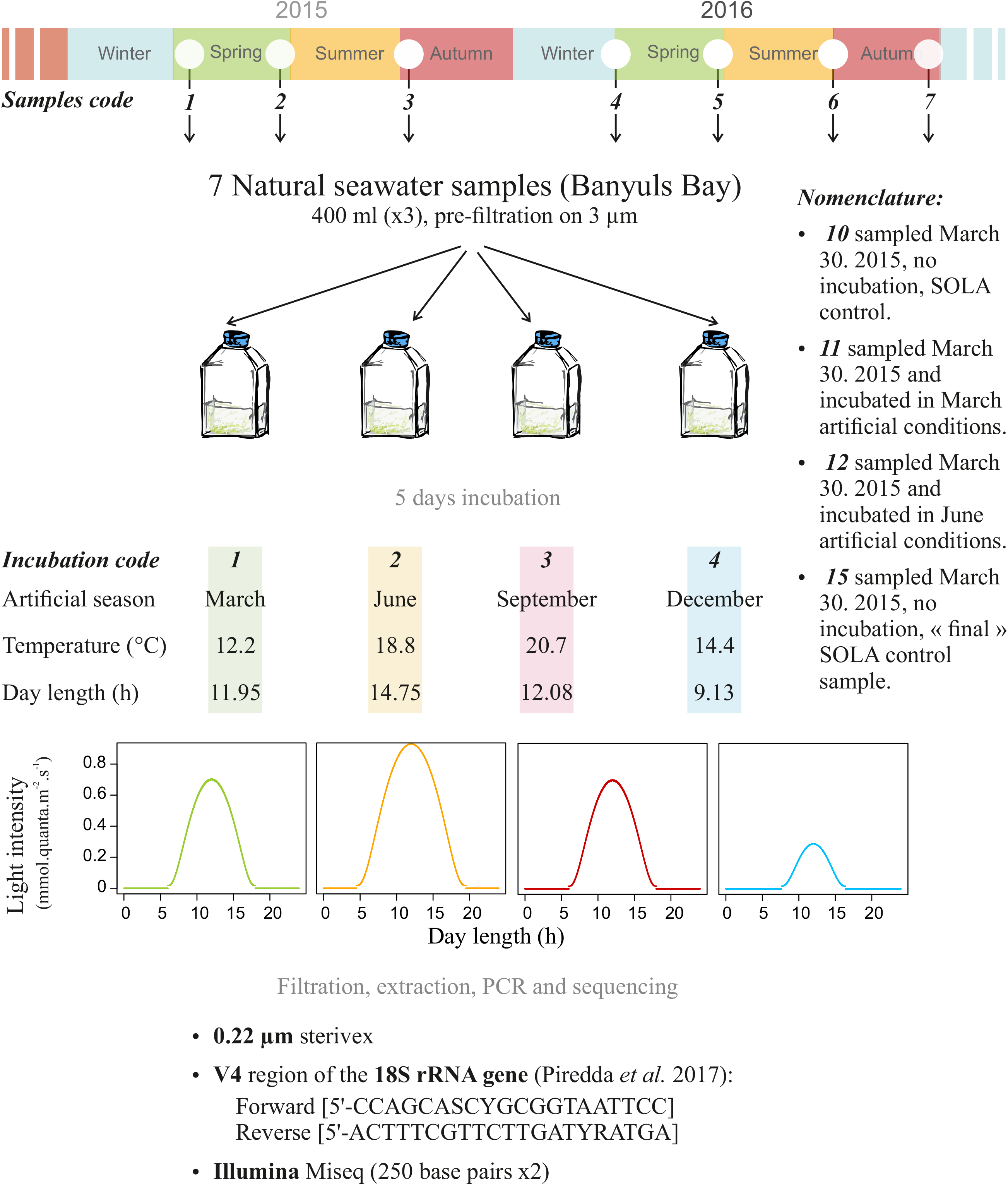
Description of the microcosm experimental procedure. Sea water was sampled 3 to 4 times a year in 2015 and 2016 at SOLA SOMLIT station off Banyuls-sur-mer (France). The 7 seawater samples were incubated in artificial conditions mimicking average seasons. Each experiment was re-named by a code combining the sampling date (1 to 7) and the incubation conditions code. Incubation codes were 0 to 5: 0 was SOLA control sample, 1 to 4 refers to incubation conditions (spring – green - 1, summer – yellow - 2, autumn – red - 3 and winter – blue - 4) and 5 refers to SOLA next control sample (4 to 10 days after initial sampling).

### DNA extraction, amplification and sequencing

DNA extracted from SOLA high-frequency time series and microcosm experiments were amplified and sequenced. DNA extractions were performed following the same protocol as Hugoni *et al* [20] and Lambert *et al* [9]. The V4 region of the 18S rRNA gene was amplified using the TAReuk primers 5’-CCAGCASCYGCGGTAATTCC (forward) and 5’-ACTTTCGTTCTTGATYRATGA (reverse) modified from Stoeck *et al* [21]. Then, paired-end Illumina sequencing Miseq (2×250 base pairs) was performed by the Genotoul platform (Toulouse, France). SOLA high-frequency time series metabarcoding datasets are available on NCBI under accession number PRJNA579489.

### Bioinformatics treatment of metabarcoding data

SOLA high-frequency time series and microcosms metabarcoding datasets were analyzed separately but using identical DADA2 bioinformatics pipeline version 1.6 [22] implemented in R v 3.6.1. The parameters were set as: trimLeft=c(20, 21), maxN=0, maxEE=c(2,2), truncQ=2 and truncLen=c(250,250) respectively c(280,230) to analyze the time series and the microcosms data. To allow comparison between the two datasets (e.g., in terms of shared Amplicon Sequence Variants (ASVs)), SOLA and microcosms ASVs and count tables were pulled together and re-processed under Mothur pipeline v1.35.1 [23] after keeping a single copy of each ASV using the *unique.seqs(*) function, ASVs were taxonomically re-classified using the Protist Ribosomal Reference Database v4.11.1 [PR^2^, 24, https://pr2-database.org/] in which several protists groups were recently curated such as Chlorophyta [25], Haptophyta [26] and dinoflagellates [27].

### Statistics and data visualization

Statistics and graphics were performed using R software version 3.6.1. Calculation of Simpson diversity index [28] and richness (Sup. Data 1), Bray-Curtis dissimilarity matrix used for Non-parametric Multi-Dimensional Scaling (NMDS, Sup. Data 2) and SIMPER [similarity percentage analysis, 29] test were performed using the *vegan* v. 2.5.6. R packages. Common R packages such as *Treemap* (e. g. used for Sup. Data 4) or *Hmisc* to plot standard deviation were also used to produce figures. All R scripts were formatted using the *Rmd* R packages and are available in supplementary data (Sup. Data 1to Sup. Data 5).

## Results

### Flow cytometry analysis of microcosms

Microcosm incubation experiments were performed in parallel to the high-frequency time series at SOLA buoy in the bay of Banyuls (42°31′N, 03°11′E, France) between January 2015 and March 2017 [30]. Microbial community from SOLA were sampled and filtered at seven dates (between spring 2015 and autumn 2016) corresponding to each of the four seasons (Supplementary table 1). Samples filtered on 3 μm were exposed in triplicates to realistic light conditions (photoperiod and light intensity) and temperature of the four seasons at the Banyuls latitude (Fig.1). After five days, photosynthetic cell abundance ranged from 290 (December 2016) to 32 000 cells.ml^−1^ (March 2015) with an average value of 6 300 cells.ml^−1^ in microcosms lacking nutrients supplementation (Supplementary Fig. 1 A) and from 540 cells.ml^−1^ (June 2015) to 100 700 cells.ml^−1^ (March 2015) with an average value of 11 730 cells.ml^−1^ in NP enriched microcosms (Supplementary Fig. 1 B). Overall, the triplicate showed the same pattern of growth even though deviations were occasionally observed (e.g., for sample 3-3 at day 5). Different patterns of picophytoplankton growth were observed between microcosms. While cell growth was observed in microcosms 1 and 7, an increase in cell number occurred only at day 5 for Microcosms 2 and 5 and a decrease of picophytoplankton cells in all triplicates was observed over the time course of microcosm 4.

### Major eukaryotic groups recovered by metabarcoding

Microcosms and environmental samples metabarcodes represented around 2 million reads, which were distributed into 2395 ASVs (Supplementary Table 2). Only ASVs represented by more than ten identical reads were taken into account. ASVs size ranged from 11 to 271 707 reads (ASV assigned to Syndiniales) with an average size of 62 reads. ASVs were assigned to the nine eukaryotic supergroups present in the PR^2^ database taxonomy [24], yet 373 ASVs were unassigned at this taxonomic level. Stramenopiles, Alveolata, Archaeplastida and Rhizaria supergroups were the four most represented in microcosm ASVs (Fig. 2 A) while Alveolata, Stramenopiles, Archaeplastida and Hacrobia were the main supergroups recovered from the SOLA environmental time series ASVs (Fig. 2 B). Reproducible seasonal patterns of ASV abundance were observed for both photosynthetic and heterotrophic (e.g. Opisthokonta and Alveolata) eukaryotes (Supplementary Fig. 2 A). Photosynthetic eukaryotes were detected primarily between late autumn and early Spring each year (Supplementary Fig. 2 B).

**Fig. 2:**
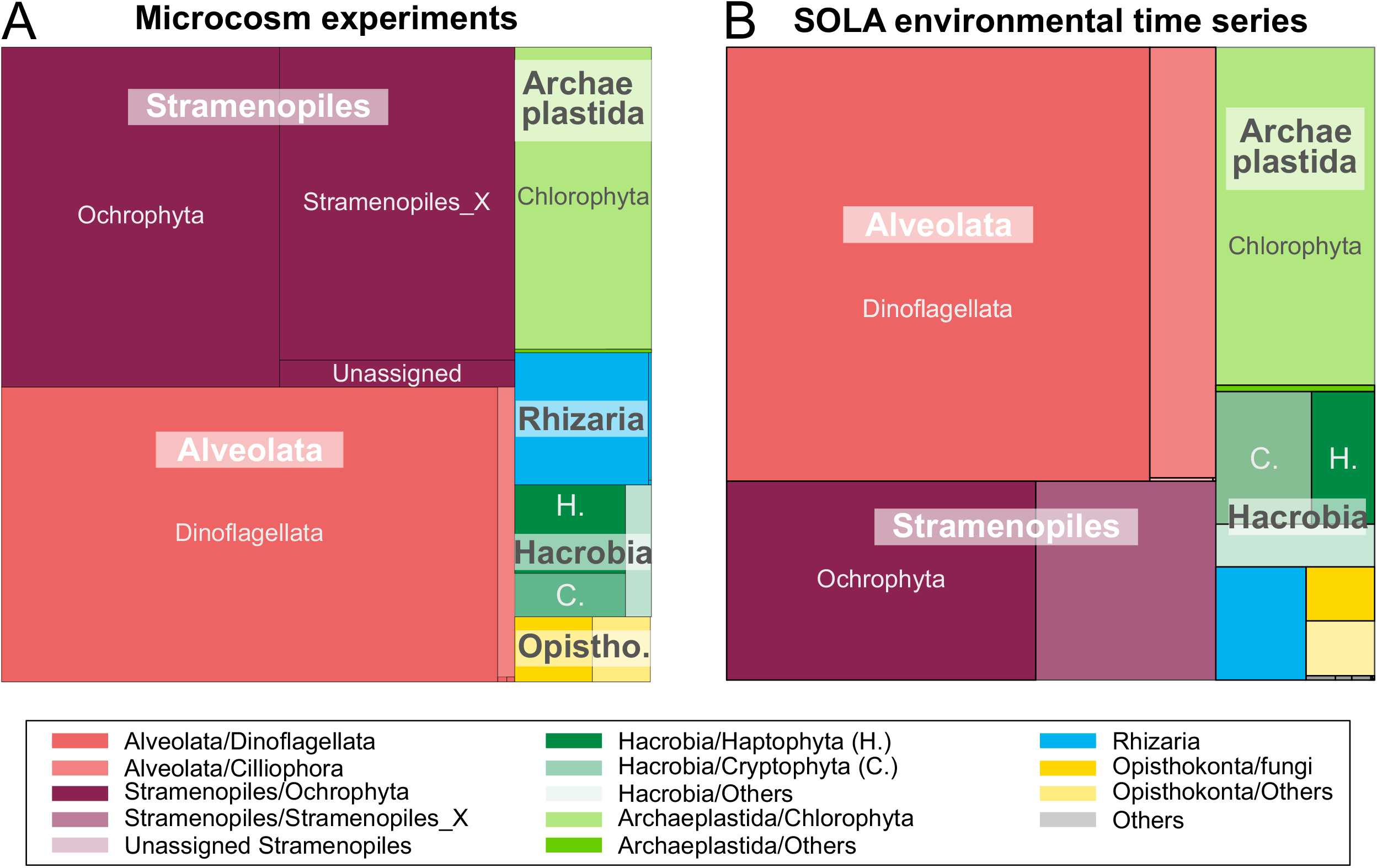
Treemaps representing the distribution of V4 region of the 18S rRNA gene metabarcodes in supergroups and major divisions of marine protists. Bold legends refer to lineages hosting photosynthetic protists. A- Global taxonomic diversity in the 56 microcosm samples, B- Global taxonomic diversity in 140 SOLA times series samples.

Stramenopiles were represented by 503 559 reads split into 671 ASVs, in which 272 791 reads (373 ASVs) were assigned to the photosynthetic Ochrophyta division (Fig. 2). In this division, reads were also assigned to the environmental clades MAST (marine Stramenopiles, 153 204 reads, 156 ASVs) and MOCH (marine Ochrophyta, 105 492 reads, 28 ASVs, [31]). In SOLA, Diatoms (Bacillaryophyta) drove Ochrophyta temporal dynamics, especially in late autumn, winter and early spring (Supplementary Fig. 3).

Alveolata were represented by 436 920 reads (assigned to 757 ASVs), in which most were assigned to dinoflagellates (422 358 reads, 628 ASVs). In SOLA, Alveolata reads were assigned to Dinophyceae, Syndiniales (especially the Dino-group I clade 1) and to a lower extent to grazers such as the Spirotricheae Ciliophora (Supplementary Fig. 2). For example, in sample 2-1, 85% of Alveolata reads corresponded to Syndiniales, which have been described as parasites of protists [32, 33].

Archaeplastida supergroup was represented by 12,612 assigned reads (137 ASVs), including Chlorophyta taxa (119 200 reads, 104 ASVs). In SOLA, Chlorophyta was the second major photosynthetic group (Dinoflagellates excluded). Dinoflagellates are commonly excluded from photosynthetic groups when using metabarcoding because only half of the species are photosynthetic. Moreover, Alveolata have large repeated genomes with numerous copies of the 18S rRNA gene, that can bias reads relative counts [34–36]. Remarkably, even with a limited number of 18S rRNA genes copies per genome, Chlorophyta numbers of reads were as high as those of Syndiniales (Supplementary Fig. 2 B) and Ochrophyta (Supplementary Fig. 2 C) and even higher in winters 2015-16. The succession of Chlorophyta classes is detailed below.

### Microcosms ecological patterns

The number of ASVs (i.e., richness) ranged from 52 in sample 1-4NP (water sampled in March 2015 and incubated in December artificial conditions with nitrate and phosphate enrichment) to 636 in sample 6-4 (seawater sampled in October 2016 and set in December artificial conditions, Supplementary Table 1). On average, 177 ASVs were recovered per sample. The Simpson diversity index ranged from 0.41 in sample 5-4NP to 0.99, with an average Simpson index of 0.87. During microcosm experiments, the diversity stayed high (between 0.8 and 1) except for June 2016 (Fig. 3), while the richness decreased in microcosm experiments compared to natural samples (Fig. 3). In general, microcosms acted as a filter, which reduced the number of ASVs without unbalancing the proportion of these ASVs in samples except for 4 June sampled microcosms and sample 6-4, which showed a higher richness but the same diversity than its relative natural counterpart samples (6-0 and 6-5). The natural samples 4-5 and 7-5 differed from other natural samples by their low Simpson’s diversity index (around 6.5) and low richness (< 100 ASVs, Fig. 3). The four June microcosm (2-1, 5-1, 5-1NP and 5-4NP) showed the lowest Simpson’s diversity index in the dataset and a low richness (< 100 ASVs, Fig. 3).

**Fig. 3:**
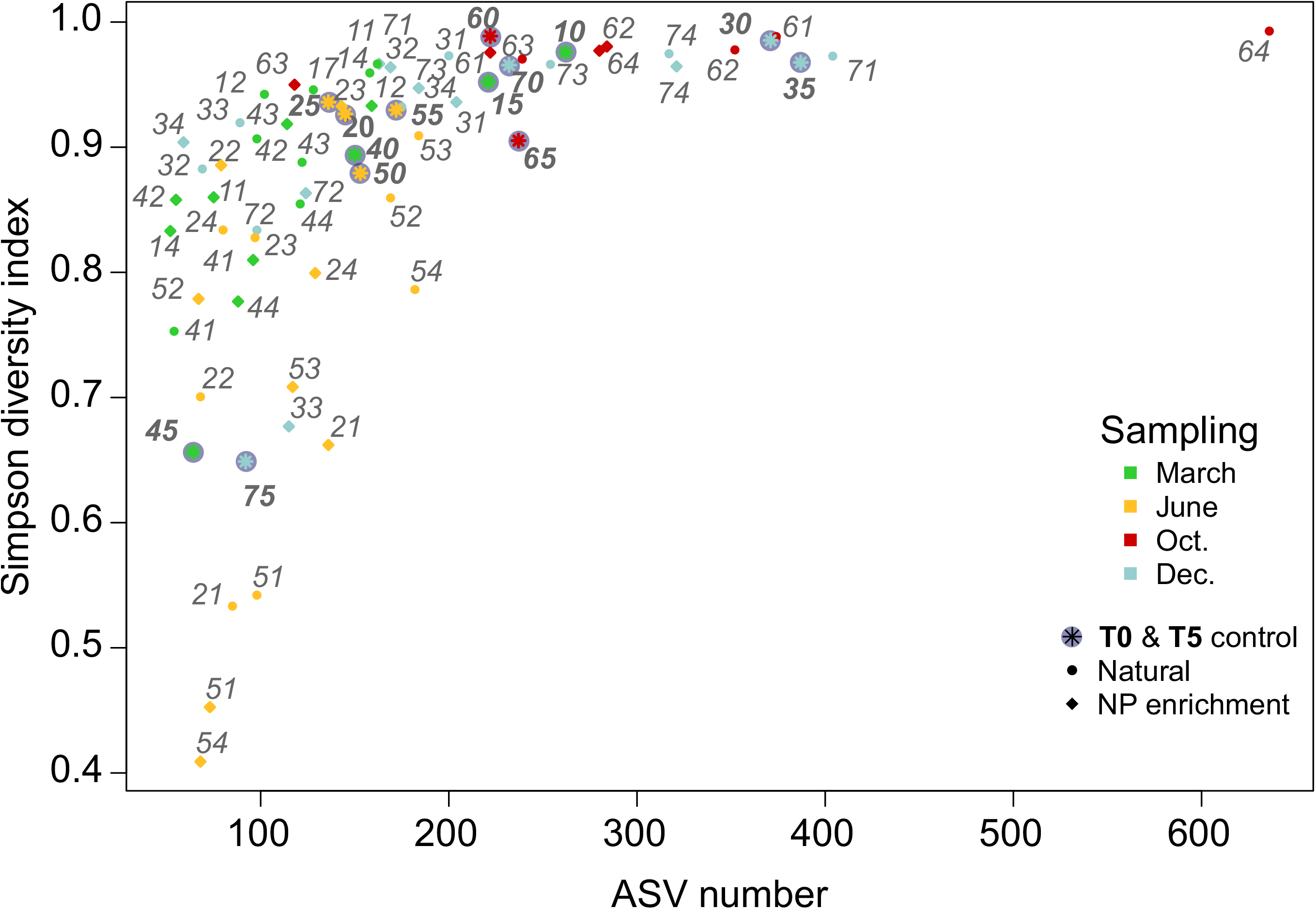
Number of ASV per microcosm samples versus the Simpson diversity index. Only ASVs represented by more than 10 reads and sample with more than 4000 sequenced reads were taken into account. Each dot and label correspond to a microcosm experiment number. Colors represent the month when the sea water was sampled and the dot shape represent either the nutriment conditions (incubation with Nitrate and Phosphate enrichment) or the natural samples from environmental time series (bold labels). Labels refer to the sampling-incubation condition codes as defined in Fig. 1 and Supplementary Table 1.

The representation of all natural and microcosms samples on an NMDS plot revealed that both natural communities and microcosm incubated samples were grouped according to sampling time rather than incubation conditions (Fig. 4). Natural samples clustered together with microcosms in June and October, March 2016 and December 2016 but were clearly separated in March 2015 (1-0 and 1-5) and December 2015 (3-0 and 3-5) indicating that changes in microbial communities had occurred in microcosm 1 and 3. Finally, environmental samples (4-0 and 4-5) as well as (7-0 and 7-5) were well separated revealing that changes in communities had occurred in the field during microcosms. Microcosms incubated with or without nitrate and phosphate (NP) enrichment clustered together, suggesting that the enrichment did not induce major changes in microbial communities. (Procrustean statistic confirmed that microcosms with and without NP enrichment were highly correlated (R^2^=0.86, Pvalue=9.9e^−5^). For this reason, only microcosms lacking NP enrichment were kept in the analysis thereafter.

**Fig. 4:**
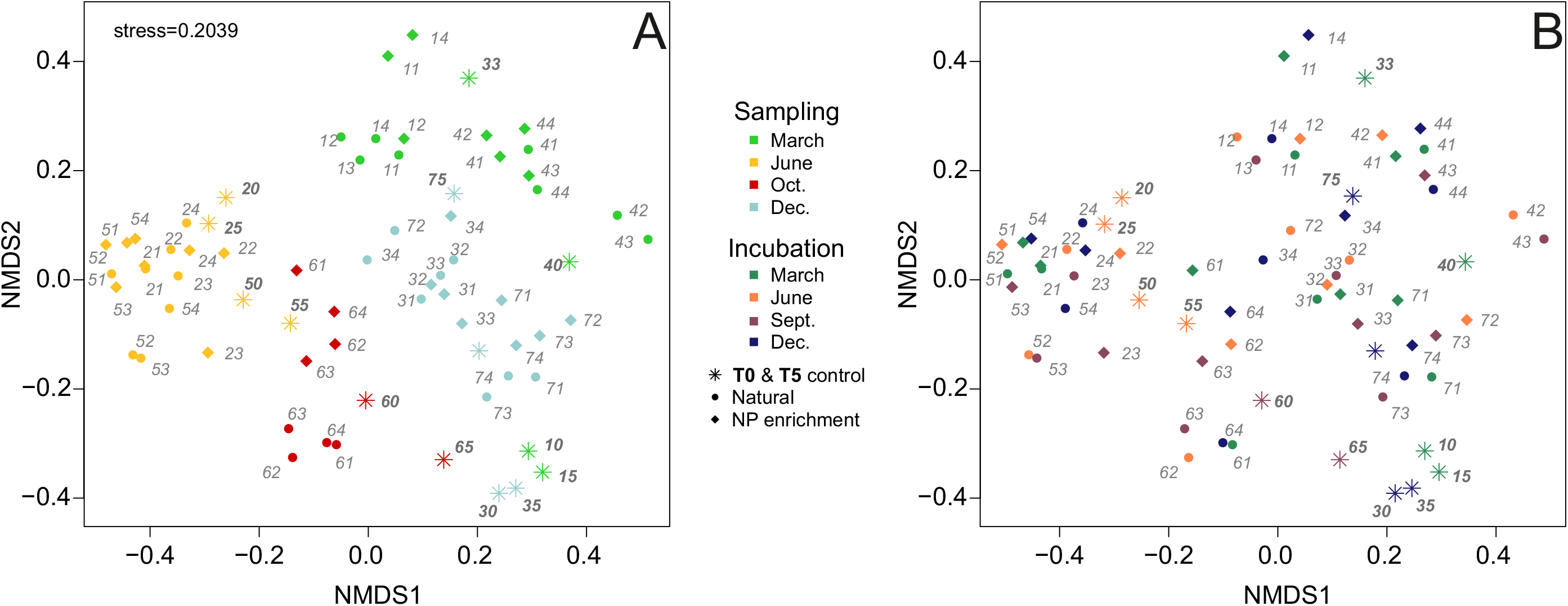
Non-parametric Multi-Dimensional Scaling (NMDS) representing microcosm experiment and natural samples protist communities. The NMDS calculation was based on a Bray-Curtis distance matrix and resulted in an acceptable stress value (0.2039). Only ASVs represented by more than 10 reads and sample with more than 4000 sequenced reads were taken into account. No environmental parameters are plotted on the NMDS graphic since none had a P-value (100 permutations) lower than 0.05. Labels refers to the sampling-incubation condition codes as defined in Fig. 1 and Supplementary Fig. 1 and the dot shape represents either the nutriment conditions (incubation with Nitrate and Phosphate enrichment) or the natural samples from environmental time series (bold labels). A-Colors represents the month when the sea water was sampled. B-Colors represent the incubation conditions (artificial month: temperature and daylight length mimicking seasons).

Most microcosm samples showed an increase in the relative abundance of Alveolata (e.g., Syndiniales dino group I clade and in a lesser extent *Gyrodinium* sp.), Opisthokonta (microcosms 1-1 to 1-4, 2-1 to 2-4, 5-2, 5-3) or Rhizaria (for example in 2-2, 2-3, 5-2, 5-3, 6-2 and 6-3 samples) relative contributions (Fig. 4, Supplementary Fig. 2). These three groups were the most represented heterotrophic protists. Concerning photosynthetic groups, several microcosms (such as 4-2, 4-3, 5-2, 5-3, 7-1 and 7-3 samples, Fig. 5) differed from the others by their major contribution of Stramenopiles, which were usually associated to warmer incubations conditions (June and September). The Chlorophyta relative contributions were higher in microcosms sampled in March and June (sample 1-2 to 1-4, 2-4, 5-4, 4-1 and 4-4 for example, Fig. 5).

**Fig. 5:**
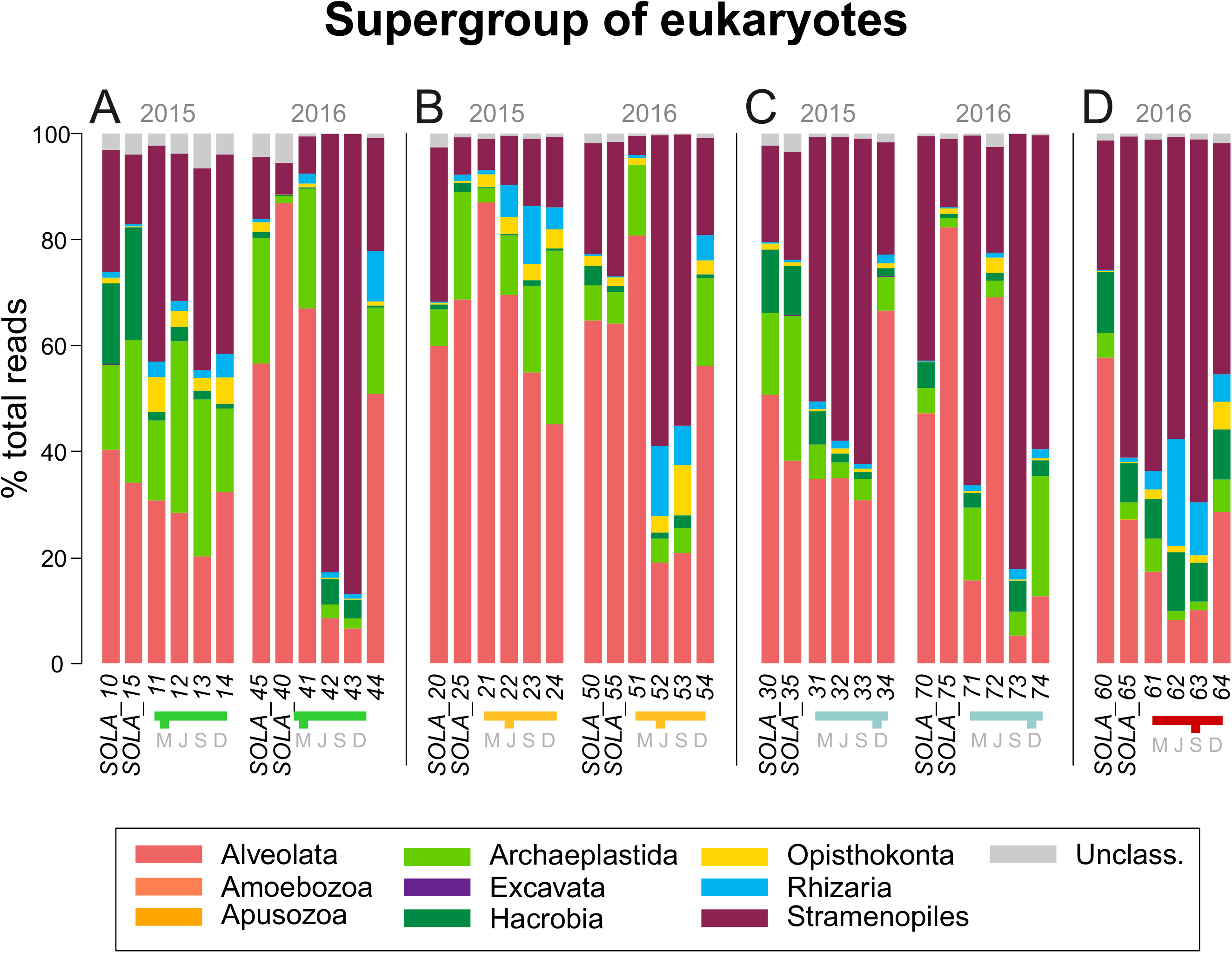
Barplot representing the percentage of reads assigned to each eukaryotic taxonomic supergroup in microcosm incubated without Nitrate and Phosphate enrichment and natural samples. Sequences assigned to Metazoans were deleted from the dataset. Colors in the barplot refer to supergroups, colors in the legend correspond to the month, when the initial sea water was sample and letters (in grey) to the incubation conditions (i.e. artificial month): March (M), June (J), September (S) and December (D). A-Natural and incubated samples from March 2015 and 2016. B-Natural and incubated samples from June 2015 and 2016. C-Natural and incubated samples from December 2015 and 2016. D-Natural and incubated samples from September 2016.

### Chlorophyta

We next focused on Chlorophyta, which provides an interesting case study of seasonal succession with ASVs belonging to this group found all year round in SOLA. In the time series, 161 ASVs were assigned to Chlorophyta, among which Mamiellophyceae (41 ASVs) and Chlorodendrophyceae (9 ASVs) dominated the green microalgae community, both showing clear seasonal patterns (Fig. 6 A). Mamiellophyceae represented 100 % of Chlorophyta reads between January and March 2015 and in 2016, and more than 80 % between October and December 2016. Chlorodendrophyceae represented up to 60 % of Chlorophyta reads from April to early September in 2015 and 2016. The number of reads assigned to other Chlorophyta classes and their relative contribution to Chlorophyta was too low to identify clear seasonal patterns. Mamiellophyceae and Chlorodendrophyceae did rarely co-occur in time (Fig. 6 A). Mamiellophyceae reads were recovered mainly in winter and early spring while Chlorodendrophyceae reads followed Mamiellophyceae peaks of abundance from late spring to late summer.

**Fig. 6:**
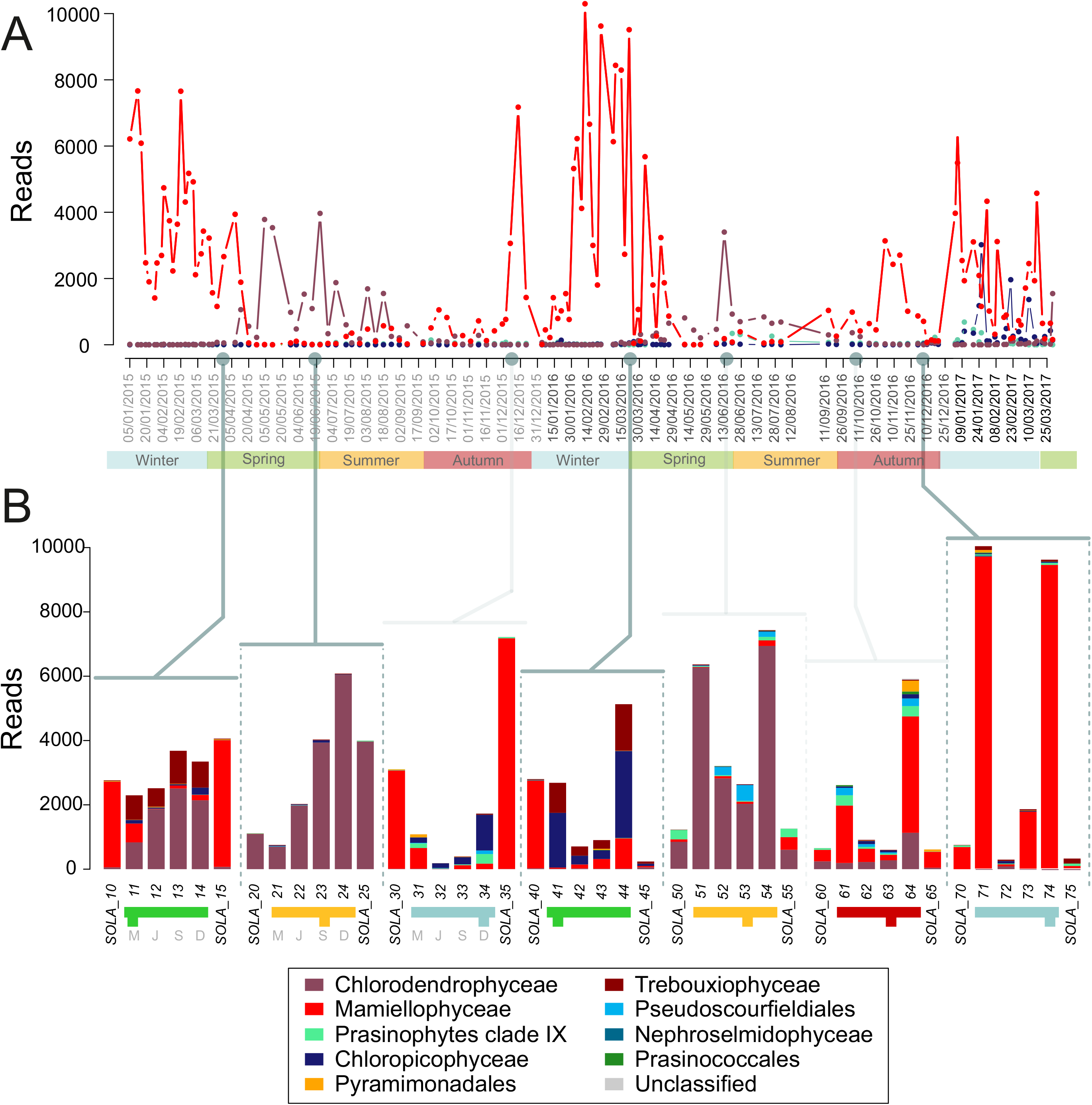
Chlorophyta classes contribution: A-Relative contribution of Chlorophyta classes (%) in the SOLA environmental high sampling frequency time series. Grey dots and lines point out the sampling date of sea water incubated for microcosm experiments. B-Barplots represent the number of reads per Chlorophyta classes in the 4 microcosm experiments and their associated natural sample. Colors in the barplot refer to Chlorophyta classes, colors in the legend correspond to the month, when the initial sea water was sample and letters (in grey) to the incubation conditions (i.e. artificial month): March (M), June (J), September (S) and December (D).

In microcosm experiments, the relative contribution of Chlorophyta usually decreased to the benefit of Alveolata or Stramenopiles reads, except in microcosm 1 (March 2015, Fig. 5). SIMPER analyses found between 2 and 4 ASVs assigned to Chlorophyta (e.g., M0005 *Bathycoccus*, M0010 *Micromonas* or M0050 Chlorodendrales) involved in the dissimilarity between samples in all incubations except for microcosm 6 (Supplementary Fig. 2). In the microcosm metabarcoding dataset, Chlorophyta reads were mainly assigned to Mamiellophyceae or Chlorodendrophyceae classes, but also to Chloropicophyceae (samples 3-2 to 3-4, 4-1 and 4-4) and Trebouxiophyceae (samples 1-1 to 1-4, 4-1 and 4-4, Fig. 6 B). Mamiellophycae dominated both incubated and natural samples in microcosms 3, 4 and 7, while Chlorodendrophyceae dominated both incubated and natural samples in microcosms 2 and 5. In microcosm experiments 1 and 4, the dominant Chlorophyta class switched between natural and incubation samples indicating that incubations profoundly modified the Chlorophyta community. Experiment 1 and 4 (March) showed respectively Chlorodendrophyceae (ASV M0050, Supplementary Table 3) and Chloropicophyceae (ASV M0037, Supplementary Table 3) as dominant Chlorophyta class, while Mamiellophyceae dominated natural samples sampled (1-0, 1-5 and 4-0, since very low reads of Chlorophyta were assigned to 4-5, Fig. 6).

Moving to the genus taxonomic level, Mamiellophyceae dominant genera in the SOLA time series were *Micromonas* and *Bathycoccus* (Supplementary Fig. 5). Across the three studied winters (2015, 2016 and 2017), *Micromonas* peaks arrived in early winter followed by *Bathycoccus* peaks in late winter or early spring (Supplementary Fig. 4 A). In microcosms, the number of reads assigned to Mamiellophyceae was low in samples from the microcosms 1, 3 and 4. Nevertheless, Simper statistic tests unveiled that *Micromonas* (ASVs M0010 and M0046) and *Bathycoccus* (M0005) contributed significantly to the dissimilarities between samples in experiments 1, 3, 4 and 7 (Supplementary Table 3). *Micromonas* reads were recovered in samples 7-1, 7-3 and 7-4, while *Bathycoccus* dominated sample 4-4 (Supplementary Fig. 4 B). Mamiellophyceae (especially *Micromonas*) were usually more abundant under low temperature, i.e. when the applied temperature in microcosms was similar or lower than the environmental temperature (i.e., in conditions 1-March and 4-December the temperature was circa 14°C).

## Discussion

### Metabarcoding versus flow cytometry

Metabarcoding of the 18S rRNA gene V4 region revealed changes in microbial community composiion in response to seasonal conditions of light and temperature in all microcosms.Metabarcoding of 18S rRNA is a non-selective but also non-quantitative method since it depends on the number of 18S rRNA gene copies in protists genomes. Alveolata and to a lesser extent in Stramenopiles supergroups, which have large genomes with numerous copies of 18S rRNA genes, are usually overestimated in metabarcodes datasets [35, 36]. In contrast, Chlorophyta have relatively small genomes and few copies of the 18S rRNA gene (e.g., 3 in *Ostreococcus tauri*) that lead to underestimating their number. Metabarcoding, therefore provides a good picture of the relative contribution of protists in global marine microbial communities [37] as well as of the diversity within specific groups such as diatoms [38] or green microalgae [11] but does not provide information on the impact of cell growth or mortality on community composition. Although less resolutive than metabarcoding, flow cytometry is a quantitative method that captures the daily change of cell numbers in microcosms over five days.

Changes in photosynthetic cell numbers, as determined by flow cytometry, were consistent between triplicates in most microcosms, although more variability was occasionally observed as seen in samples 7-2 and 3-3 (Supplementary Fig. 1). Several factors could account for this variability. Though a filtration protocol was used to remove nano-phytoplankton containing predators, some larger-sized grazers of picophytoplankton can sometimes pass through the 3 μm filter and induce disturbances in microcosms.

Flow cytometry data showed season dependent growth patterns that could be potentially explained by the initial density of photosynthetic cells in microbial communities. In winter, low photosynthetic cells density allows an exponential growth pattern in each experiment, while in June and March, photosynthetic communities should face both higher competition between microalgae and interactions such as parasitism or grazing, which generally occurs at higher photosynthetic cells density [7, 14].

### Effects of five days incubation on microbial communities

The five-day incubations affected the different taxa differently, as illustrated for Hacrobia a division which comprises Cryptophyta and Haptophyta. For example, the contribution of Cryptophyta drastically decreased in all samples of microcosms 1 and 3 (March 2015 and December 2015), respectively, while natural samples showed a relatively high contribution of Cryptophyta reads (Fig. 5). In microcosm 6 (October 2016), however, the relative contribution of Haptophyta remains stable between natural communities and microcosms. These observations suggest that, unlike Haptophyta, Cryptophyta from the Banyuls Bay may be too difficult to grow or maintain in microcosms.

Stramenopiles was the division that showed the highest differences between microcosms and environmental samples (T5). Its relative contribution showed massive increase in several microcosms (such as 4-2, 4-3, 5-2, 5-3, 3-1, 3-2, 3-3, 7-1, 7-3, 7-4 and experiment 6 samples, Fig. 5). This increase in relative contribution should be interpreted in the light of flow cytometry data on day 5. In microcosms 4 (March 2016), we observed a dramatic decrease in photosynthetic cell number (from 10 000 events to 2000), suggesting that the relative increase of Stramenopiles may result from cell mortality in other divisions, such as Archeaplastida. In microcosm 7 (December 2016) in contrast, the global increase in photosynthetic cells observed by flow cytometry suggested that this increase could correspond to diatoms bloom (Ochrophyta, Simper test,Supplementary Table 3). Diatoms occurred in several size fractions [39, 40] and recovering blooms of diatoms in picoeukaryotes microcosms were expected as they are an important phytoplanktonic organism in Mediterranean Sea waters and were already described to follow clear seasonal patterns [41, 42].

When Stramenopiles were less abundant, the natural communities and microcosms were often dominated by Alveolata, particularly Syndiniales [which contain known paratisoids, 32, 33] were clearly more abundant in microcosms than in environmental samples. In general, the higher relative contribution of heterotrophic protists and putative parasites, such as Syndiniales and *Labyrinthule* in microcosms suggest that the incubation process may enhance infection processes. In addition, the presence of bacterial grazers such as Choenoflagellates, or the ciliate, as *Minoresa minuta* yet [43] evoked that post-bloom conditions were met after five days of incubation days with a decrease of the photosynthetic contribution to the benefit of heterotrophs in several microcosms. The effect of “bottle enclosure” in small volume may be responsible for speeding up biological processes and unbalancing the autotrophic to heterotrophic ratio in microcosms [16, 44].

### Effects of light and temperature on microbial community composition in microcosms

It is usually challenging to discriminate between light and temperature effects on microbial communities since these two parameters co-vary in the environment. March and September have similar day length and light conditions among the four seasonal conditions but have an 8°C temperature difference. Comparing microcosms conditions of March and September allows thus, to assess the respective effects of light and temperature. Incubations under these two months’ conditions led to dramatic changes in community compositions, highlighting the importance of temperature in shaping microbial communities. In particular, Stramenopiles’ relative contribution increased markedly in September warm condition but not in March cold condition as seen in microcosms. Stramenopiles contribution was also high under the June warm condition, confirming the temperature effect on Stramenopiles. Recent microcosm studies have emphasized the putative role of diatoms in a warming ocean, consistent with Stramenopiles exceeding natural relative contribution in the warmest incubated conditions [18]. However, it should be noted that in microcosms 3 and 6, the Stramenopiles contribution was high in a majority of conditions.

### Influence of initial communities

Overall, the five-day incubation did not change the protist community enough to switch the incubated community to a community representative of the applied seasonal condition in most microcosms (Fig. 4).. The inter-annual variations between 2015 and 2016 could explain most of the differences seen between microcosms sampled during similar seasons. In general, microcosms and natural samples 2015 and 2016 clustered together (Fig. 4), except for natural samples from microcosms 1 and 3 (March and December 2015). This suggested that winter 2015 was different from winter 2016. The environmental parameters, especially temperature and Chl, measured at the SOLA SOMLIT Station were similar between March 2015 and 2016 (around 12°C and 0.8-1.7 μg/L). Inter-annual variability was driven mainly by dinoflagellates blooms, which could account for the observed inter-annual differences in March and December between 2015 and 2016 (Supplementary Table 3). Differences between natural samples and microcosms dynamics were also observed in March 2015 (microcosm 1) in which the incubation condition was sufficient to move the march 2015 natural community towards a “2016 like” community. The increase of temperature in 2015 induced a dinoflagellates bloom (e.g. *Gyrodinium* sp.) even though dinoflagellates were not blooming in natural samples. The blooming dynamics allow photosynthetic protists to rapidly dominate communities when environmental conditions became favorable. Blooms occurring in microcosms suggest that microalgae had encountered both favorable environmental conditions and out-competed other protists.

The dynamics of natural communities also seem to affect the outcome of microcosms. In microcosms 4 and 5 sampled before a bloom (Supplementary Fig. 2 B, Supplementary Fig. 3) Chrysiophyceae gold microalgae bloomed only in incubation conditions of warm temperature (Fig. 5, Supplementary Table 3). On the contrary, in microcosms 3 (December), 6 (October 2016) and 7 (December 2016) sampled at the beginning or during a diatom bloom, Stramenopiles bloomed under all applied seasonal conditions even at a lower temperature (Fig. 5, Supplementary Fig. 1). These results lead to the idea of a “community inertia” for blooming species such as diatoms: When the blooming dynamics was already triggered, this dynamics continued independently of the environmental parameters applied. In contrast, under pre-bloom conditions, the environmental parameters strongly influenced the communities in microcosms.

### Temporal succession of Chlorophyta

Chlorophyta, in particular Mamiellophyceae, provide an interesting case study since their contribution is major in coastal waters [25, 45]. *Micromonas* and *Bathycoccus* peak in winter in the northern Mediterranean Sea where they can represent more than 50% of picoeukaryotes reads in winter with strong seasonal rhythms [6, 9]. The presence of seasonal Chlorodendrophyceae was only recently confirmed using the V4 region of the 18S rRNA gene metabarcoding technique [10, 11]. Chlorodendrophyceae is an abundant yet poorly documented Chlorophyta class and the major ASVs assigned to Chlorodendrophyceae in SOLA natural samples and microcosms could be assigned only at the order level (Chlorodendrales) and to uncultured organisms (i.e., environmental sequences). SOLA Chlorodendrophyceae seasonal pattern (Fig. 5) was similar to this of *Tetraselmis wettsteinii* [46], a Chlorodendrophyceae species which was shown to form massive green blooms in the Bay of Naples in late spring and summer [47]. Since no reference sequence is available for *T. wettsteinii*, it is not possible to decipher if the abundant Chlorodendrophyceae ASV from the Banyuls Bay corresponds to this species.

Interestingly, Mamiellophyceae did not always dominate winter microcosm experiments (March and December, microcosms 1, 3, 4 and 7, Fig. 6). As discussed above, the community dynamics in the field may have prevented the growth of Mamiellophyceae even under favorable conditions in microcosms. Microcosms on pre-bloom communities (September) led to an increased contribution of *Micromonas* under the winter/ autumn incubation conditions (March, December) even though *Micromonas* ASV contributions were low in September natural communities. Although documented temperature preferences of *Micromonas* Mediterranean strains range between 25 and 30°C [48], we observe that *Micromonas* blooms around 14°C in natural communities and that winter/autumn temperatures and light conditions were more favorable to *Micromonas* in microcosms. The differences in temperature preferenda of *Micromonas* between culture strains and natural communities suggest factors other than temperature such as biotic interactions. Strong light especially may be inhibiting the growth of Mamiellophyceae in microcosms since we never observed Mamiellophyceae in June incubation conditions (similar temperature as in September incubation). Biotic interactions may also influence *Micromonas’* ecological niche, such as parasitism and competition.

Experiments 1, 4 and 5 were particularly interesting to investigate the temporal patterns of Chlorophyta that occurred in SOLA natural communities (Fig. 6). Mamiellophyceae, which dominated Chlorophyta in natural communities of March 2015, were still detected together with Chlorodendrophyceae under the low temperature incubation (12.2°C) but Chlorodendrophyceae replaced them in the other microcosms of March 2015. In March 2016, a switch between Mamiellophyceae and Chloropicophyceae occurred under low temperature incubations. These results suggest that (i) the succession of Mamiellophyceae to Chlorodendrophyceae in microcosms reflect their succession in the field and (ii) the switch from Mamiellophyceae to Chlorodendrophyceae occurs independently of temperature and light conditions and is mostly influenced by ecological processes such as biotic interactions. Reads assigned to several grazers such as ciliates or parasites such as Syndiniales were found at that time, but unfortunately no ecological relationships are known between Mamiellophyceae and specific heterotrophic protists. Furthermore viruses (which were not followed in this study) may significantly affect microalgal populations and ecological interactions [7, 49].

## Conclusion

Our combined approach of microbial diversity monitoring at SOLA together with microcosms simulating seasonal conditions of light and temperature unveiled several aspects of the regulation of phytoplankton seasonality by light and temperature: (1) overall a five days incubation was not sufficient to completely reshape the initial microbial communities, (2) the *in situ* dynamics of phytoplankton starting communities modulated the impact of applied seasonal conditions in microcosms as seen for Diatoms or Chlorodendrophyceae in bloom or post-bloom conditions. (3) In pre-bloom conditions, phytoplanktonic species were the most sensitive to light and temperature conditions. Diatoms were favored by higher temperature independently of light, while Mamiellophyceae increased under lower temperatures and lower light intensities (December and March). Altogether, our results suggest that light and temperature seasonal conditions play an importantrole in regulating phytoplankton in pre-bloom conditions and biotic interactions may be preponderant in bloom and post-bloom conditions.

## Supporting information

Microcosms experimental conditions.

Set of eight tables summarizing pairwise simper statistical test: the composition of each of the 6 samples belonging to the same microcosm experiment

## Acknowledgments

We would like to thank the captain and the crew of the RV ‘Nereis II’ for collecting the samples and the Service d’Observation”, particularly Eric Maria and Paul Labatut, for processing of the samples. We are also grateful to Urania Christaki for constructive exchanges during manuscript writing, Pierre Galand for providing advice in statistical analyses and Adam Monier for constructive reading of the manuscript. This work was supported by the French Agence Nationale de la Recherche through the Photo-Phyto project (ANR-14-CE02-0018) to FYB.

## Supplementary Information

**Supplementary Table 1:**Microcosms experimental conditions. ‘Sample’ was the original sample names, that were translated into a two digit code ‘Code’. Sample names starting by ‘SOLA_’ are natural control samples extracted from the environmental time series. ‘Sequencing Date’ refers to the date of sea water filtration after 5 days incubations to prepare DNA extraction. ‘Artificial month’ refers to the light and temperatures incubation conditions: Daylight duration ranged from 9.13 hours in December to 14.75 hours in June, with maximum light intensity between 0.3 and 0.9 mmol.quanta.m^−2^.s^−1^ corresponding to the 3 meter depth of sampling at SOLA. Applied temperature corresponded to the average temperature of the sea at SOLA recorded between 2007 and 2014: in March (12.2 °C), June (18.8 °C), September (20.7 °C) and December (14.4 °C). The nutrients enrichment are coded as a Boolean variable, T-5 μM of NaNO_3_ and 0.2 μM of Na_2_HPO_4_ (SIGMA Aldrich) were added to microcosms (NP microcosms) versus F-no nutriments supplementation. All microcosms were sampled at the surface.

**Supplementary Table 2**: ASVs table of microcosm experiments and control natural samples: PR^2^ taxonomy and number of reads per samples.

**Supplementary Table 3:**Set of eight tables summarizing pairwise simper statistical test: the composition of each of the 6 samples belonging to the same microcosm experiment (microcosms 1 to 7) was compared as well as the March natural samples (1-0, 1-5, 4-0, 4-5). The taxonomical assignation of the ASVs, which influenced the most the dissimilarities between microcosms, are listed next to each table and cumulative dissimilarity % value represented by the 4 top ASVs is mentioned inside the tables.

**Supplementary Figure 1:**
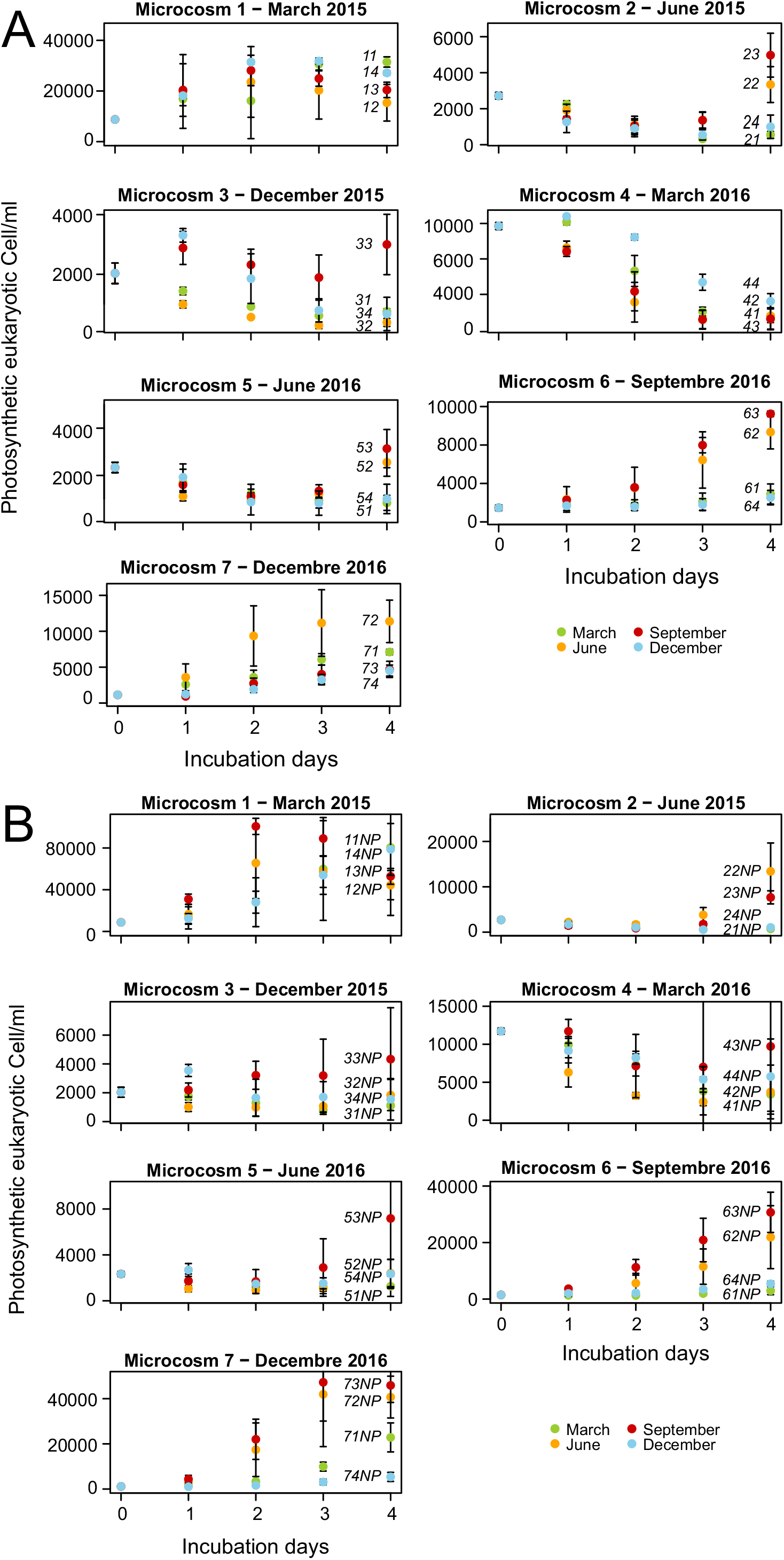
Evolution of eukaryotic photosynthetic cell numbers (pico- and nano-phytoplankton communities) during microcosm experiments incubation followed by flow cytometry measurements. The colors of the dots refers to incubation condition and the label corresponds to sampling-incubation condition codes as defined in Fig. 1 and Supplementary Table 1 **Supplementary Information**

**Supplementary Table 1**.**A**- Microcosms incubated without nutrients supplementation. **B**- Microcosms experiments enriched with nitrate and phosphate.

**Supplementary Figure 2:**
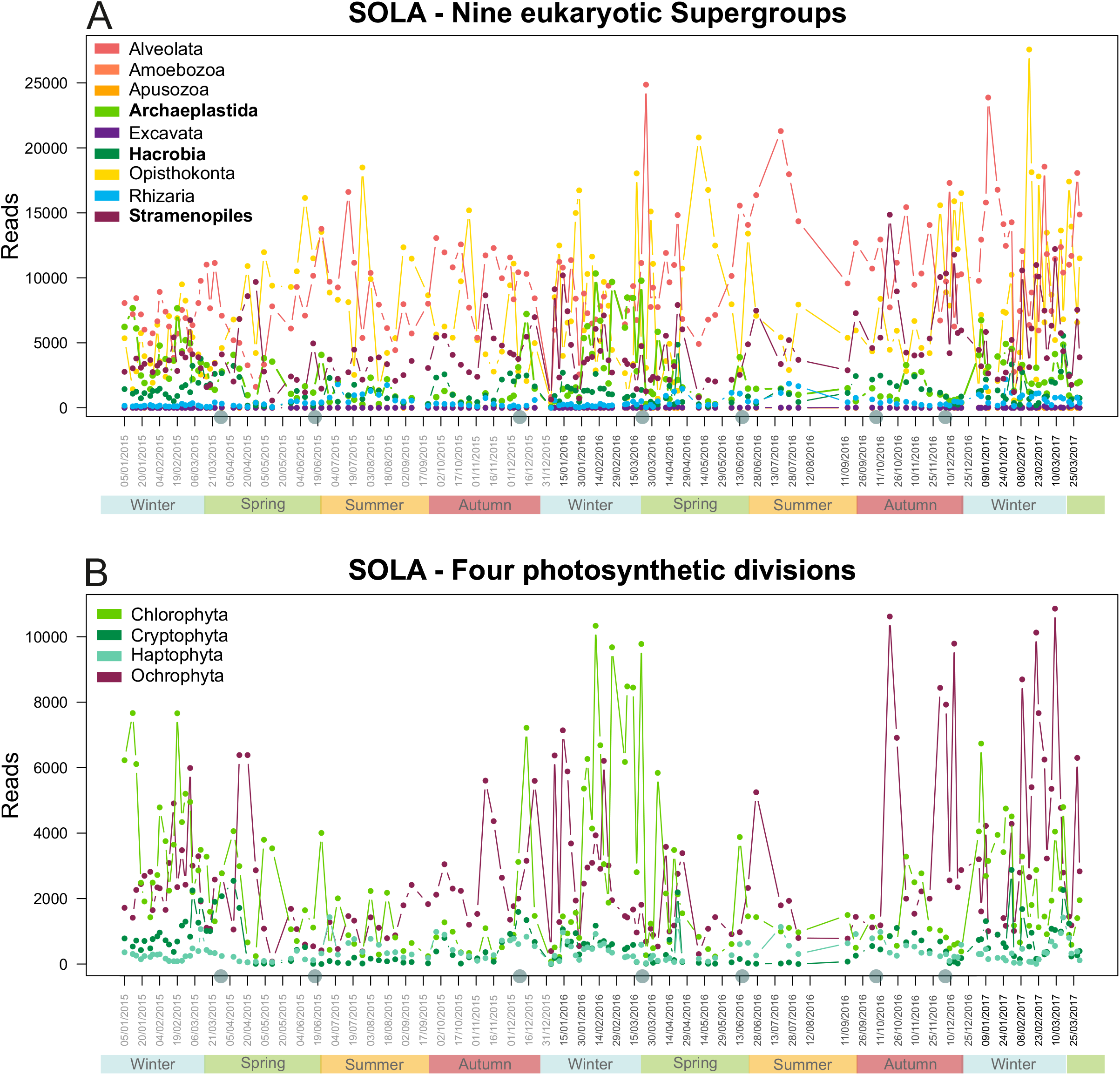
Seasonal patterns of reads assigned to protists taxonomic groups in SOLA environmental high-frequency time series. Grey dots correspond to the sampling dates of sea water for microcosms experiments. **A**- Pattern of abundance of the V4 metabarcodes assigned to the nine eukaryotic supergroups. Metabarcodes assigned to Metazoan were not deleted from the Opisthokonta. Photosynthetic supergroups are in bold in the legend. **B**- Pattern of abundance of the numbers of the V4 metabarcodes assigned to the four major photosynthetic divisions (dinoflagellates excluded).

**Supplementary Figure 3:**
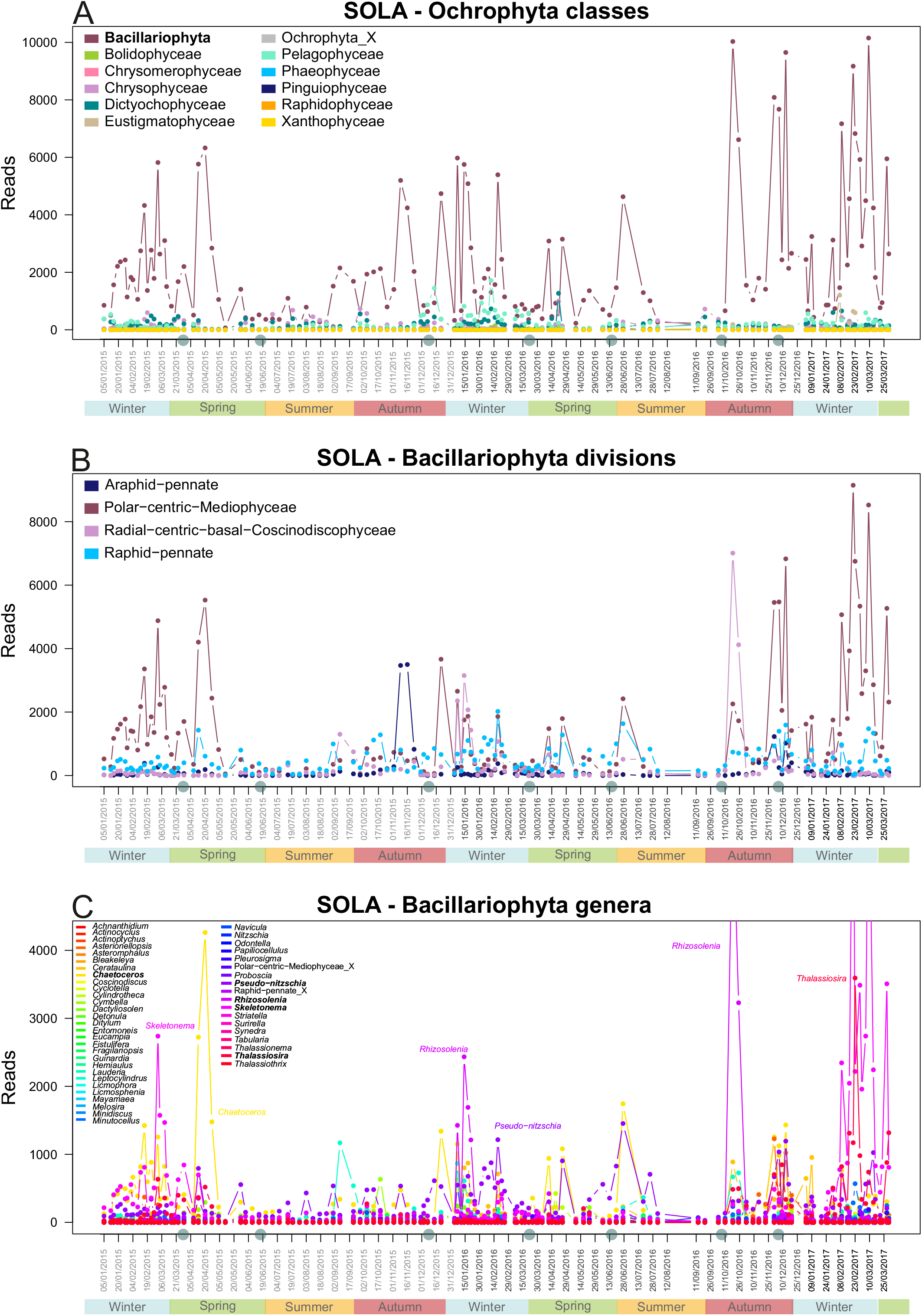
Seasonal patterns of reads assigned to Ochrophyta microalgae (photosynthetic division of Stramenopiles) in SOLA environmental high-frequency time series. Grey dots correspond to the sampling dates of sea water for microcosms experiments. **A**- Pattern of abundance of the V4 metabarcodes assigned to Ochrophyta classes. **B**- Pattern of abundance of V4 metabarcodes assigned to Bacillariophyta (Ochrophyta) families, **C**- Pattern of abundance of the numbers of the V4 metabarcodes assigned to the Bacillariophyta genera. The most abundant genera are in bold in the legend.

**Supplementary Figure 4:**
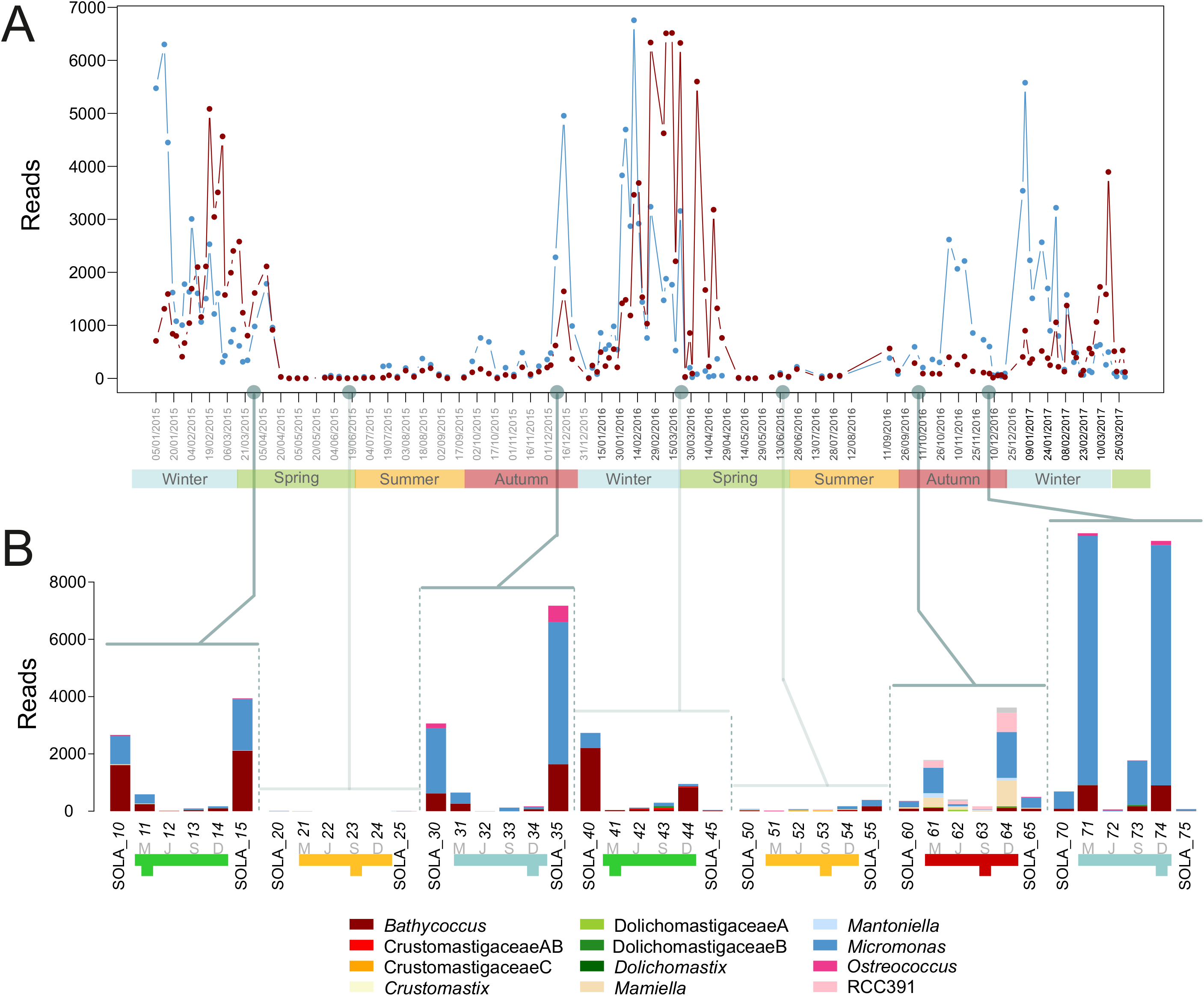
Mamiellophyceae genera contribution in SOLA environmental time series and microcosms. **A**- Seasonal patterns of reads assigned to Mamiellophyceae genera at SOLA. Note that only the two most abundant genera *Micromonas* and *Bathycoccus* can be seen on the graph. Grey dots correspond to the sampling dates of sea water for microcosms experiments. **B**- Barplot representing the number of reads per Mamiellophyceae genera in 4 microcosm experiments and their associated natural sample. Colors in the barplot refer to Mamiellophyceae genera, colors in the legend correspond to the month, when the initial sea water was sample and letters (in grey) to the incubation conditions (i.e. artificial month): March (M), June (J), September (S) and December (D).

**Supplementary Data 1:**Script used to compute and plot ASVs richness versus Simpson’s diversity index (Fig. 2) as *Rmarkdown.html* file.

**Supplementary Data 2:**Script used to compute and plot (Fig. 3) as *Rmarkdown.html* file.

**Supplementary Data 3:**Script used to compute and draw barplots (Fig. 4, Fig. 5B and Fig. S 5B) as *Rmarkdown.html* file.

**Supplementary Data 3:**Script used to compute and plot Treemap (Supplementary figure 2) as *Rmarkdown.html* file.

**Supplementary Data 5:**Script used to plot Fig. 5A, Supplementary figures 3 to 6 as *Rmarkdown.html* file.

